# Metabolism is correlative not causative for age-related auditory decline in an insect model

**DOI:** 10.1101/2022.12.20.520730

**Authors:** Thomas T Austin, Christian Thomas, Lewis Clifton, Alix Blockley, Ben Warren

**Author notes:** Lead and corresponding author: Ben Warren.

## Abstract

Aging is due to a complex decline of multiple biological processes. Some of the causes include oxidative damage, mitochondrial and proteostatic dysfunction, and DNA damage. The result is that as biological systems age their performance deteriorates. This age-related decline is well quantified, and experienced, for human hearing and is presumed to be due to a decrease in the ear’s metabolism – specifically a decrease in ability to maintain an electrochemical gradient, the endocochlear potential. However, direct measurements of metabolism across a lifespan in an auditory system are lacking. Even if metabolism does decrease with age, the question remains is it a cause of age-related auditory decline or simply correlative? All auditory systems across the animal kingdom share functional principles including ion pumping cells, auditory receptors, spiking auditory nerves and multiple supporting cells. Therefore, we used an insect, the desert locust, *Schistocerca gregaria*, as a physiologically versatile model to understand how cellular metabolism correlates with age and impacts on age-related auditory decline. We found that although metabolism correlates with age-related auditory decline it is not causative.

## Introduction: Metabolic basis of age related auditory decline - correlative or causative?

As we age cellular metabolism declines and we succumb to age-related hearing loss. This has led to the hypothesis that metabolic decline is, in part, responsible for age-related hearing loss (Vaden et al., 2017, 2022). However, both ageing and hearing loss have complex multicellular and multifaceted cellular bases. Causes of cellular ageing include: decreased proteostasis, epigenetic regulation, DNA damage, increased inflammation, mitochondrial dysfunction and decreased cellular genesis (McHugh and Gil, 2018). Age-related hearing loss has been attributed to loss of auditory receptors (Coleman, 1976; Keithley and Feldman, 1982; Bhattacharyya and Dayal, 1985; Li and Hultcrantz, 1994)), their synapses (Kujawa and Liberman, 2006, 2009) degradation of the auditory nerve (Xing et al., 2012; Altschuler et al., 2015; Panganiban et al., 2022), as well as deterioration in supporting cells that establish electrochemical gradients (Cable et al., 1993; Ohlemiller, 2009).

Hearing loss has been historically categorized as sensory, neural and metabolic and is thought to be due to apoptosis of receptor hair cells, spiral ganglion neurons or deterioration of the stria vascularis respectively (Schunkecht, 1993). Stria vascularis deterioration was thought to be the dominant cause of age-related hearing loss as it is responsible for maintaining the endocochlea potential responsible for efficient transduction and amplification. Recent analysis has spread doubt on this interpretation in humans. Although stria vascularis atrophy is the most consistent finding, (Schunkecht, 1993) in older cochleae it is the damage of hair cells that best predicts hearing function (Wu et al., 2020). In addition, some mouse models do not show a decrease of the endocohlear potential (Liu et al., 2022). Experimental approaches are diving deeper into the cellular mechanisms and uncovering subtle age-related deterioration. This work suggests that age-dependant deterioration of cellular function, without apoptosis, could explain at least some age-dependant decline in auditory function (Blockley et al., 2022; Jeng et al., 2021; Keder et al., 2020).

Four out of seven ARHL-susceptible mouse models have an age-dependant decrease in the endocochlea potential (Ohlemiller, 2009). Separate from the lateral wall, in the organ of Corti, optical measurements of a key metabolic enzyme Nicotinamide adenine dinucleitide (NAD) show a reduction of its expression (Okur et al., 2022) and function (Majumder et al., 2019). Reduction by NAD gives a good indication of metabolism as two NAD molecules are oxidized in the glycolytic pathway, three in each Kreb’s cycle and one in the electron transport chain. There is a clear precedent for a reduction in metabolism causing age-related hearing loss (Schmiedt et al., 2002), but a lack of direct measurement of metabolism auditory tissue and a lack of age-dependant correlation in natural aging models means this is far from proven. An alternative hypothesis is that such age-related changes are due to accumulative age-acquired damage as opposed to the current metabolic state of the ear.

Insects are emerging as a model to understand basic principles of hearing loss including the genetic orchestration (Keder et al., 2020), molecular homeostasis (Boyd-Gibbons et al., 2021) biomechanics (Keder et al., 2020) and physiology (Warren et al., 2021; Blockley et al., 2022). Insects, like all other animals with ears manifest an age-dependant auditory decline (Keder et al., 2020; Blockley et al., 2022) with homologous cell and cellular processes responsible, compared to established mammalian models. These similarities include decline in the function of the transduction machinery (Keder et al., 2020), auditory receptor loss and auditory nerve deterioration (Blockley et al., 2022). It is becoming increasingly clear that age-related hearing loss is due to a combination of age-related deficits in diverse cells. All auditory organs – from mammalian to invertebrate - share metabolically demanding physiological principles of function with specialised ion pumping cells maintaining a potential across receptor cells and a spiking auditory nerve to carry auditory information to the central nervous system. In addition, metabolism is as ancient as life itself and evolved over 3 billion years ago, 2.8 billion years before the evolutionary split between invertebrates and vertebrates. On these bases we justify using a high-throughput and physiologically-malleable insect model, the tympanal ear of the desert locust. With the locusts high-throughput approach we measured the metabolic rate of the Locusts’ auditory Müller’s organ and the locust’s physiological malleability enabled manipulations of ear metabolism to test if the metabolic state of the ear determines the auditory function; in other words to test if the presumed decrease in metabolism is causative for age-related auditory decline.

## Results

### Age-dependant decrease in Müller’s organ metabolism

To test if metabolism decreases in an age-dependant manner we extracted Müller’s organs from locusts of known ages post their final moult. There was a clear decline in the rate of metabolism across all ages (Figure 1A) (LM t_(269)_=−9.2, p<1×10^−16^). However, if we split the data into two halves, of the lifespan measured, we find a decline between 0-16 days post final moult (LM: t_(145)_=−4.54, p= 0.0.000009; Cohen’s d=0.902 (comparing days 0-5 with days 30-35)) but no decline in metabolism between the ages of 20-34 days post final moult (LM t_(107)_=− 0.039, p=0.969). Thus, the rate of metabolic decline decreases and levels out with an increase in age. We used three blockers of the electron transport chain: sodium azide (1 mM), antimycin (0.5 μM), rotenone (0.5 μM) for a positive control for our metabolic assay and found a decrease in metabolic output (Figure 1Bi, LM: t_(30)_=−4.633, p=0.0000656; Cohen’s d=1.637). To assess the metabolic involvement of action potential generation and auditory neuron function we used TTX and TEA to block voltage-gated sodium and potassium channels and pymetrozine to silence auditory neurons. Both pharmacological interventions resulted in no change in the rate of metabolism (Figure 1Bii, TTX&TEA: LM: t_(63)_=−0.273, p=0.786 and Figure 1Biii, pymetrozine: LM: t_(30)_=0.929, P=0.36).

**Figure 1.**
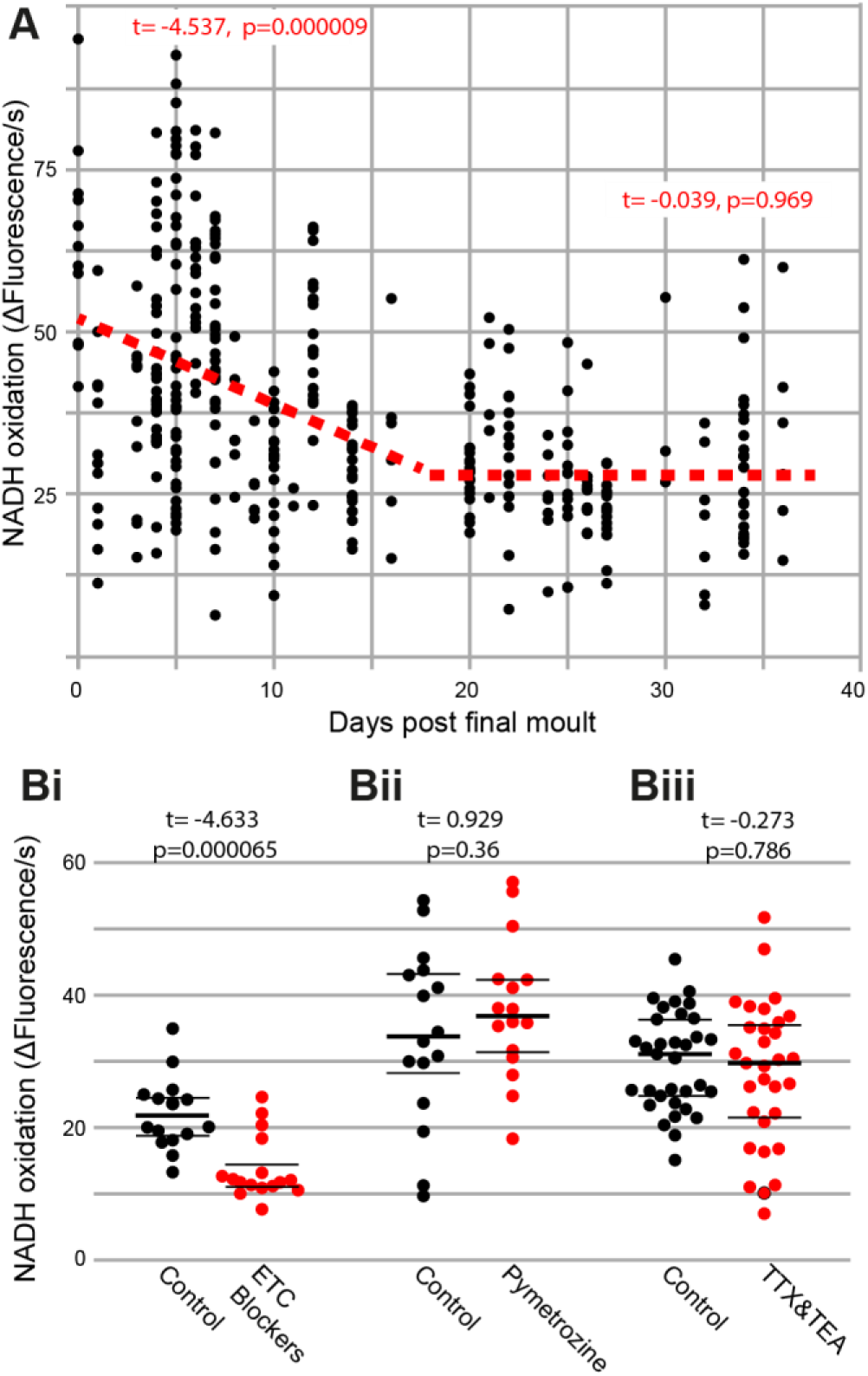
Metabolic rate decreases with age and is not effected by interfering with auditory neurons. **A**. The rate of reduction of resazurin through NADH oxidation was measured fluorometrically in two ears from individual locusts as a function of their age post last moult. The addition of electron transport chain blockers **Bi**, auditory neuron-specific insecticide pymetrozine **Bii** and sodium- and potassium-voltage gated ion channel blockers **Biii** (TTX and TEA) on the rate of resazurin reduction.

### Unchanged spontaneous auditory nerve activity with age

We recorded electrical potentials from the auditory nerve using hook electrodes from locusts 10 days to 34 days post their final moult (Fig. 2A, Bi, Bii). We either used a threshold of 100 μV to count spikes (Fig 2Biii) or calculated the standard deviation of the auditory nerve spontaneous electrical activity (Fig. 2Biv). The spontaneous spike rate did not change as a function of age (Fig. 2C, t_(127)_=0.85, p=0.397). The total electrical activity also did not change as a function of age (Fig. 2D, t_(119)_=−0.302, p=0.763).

**Figure 2.**
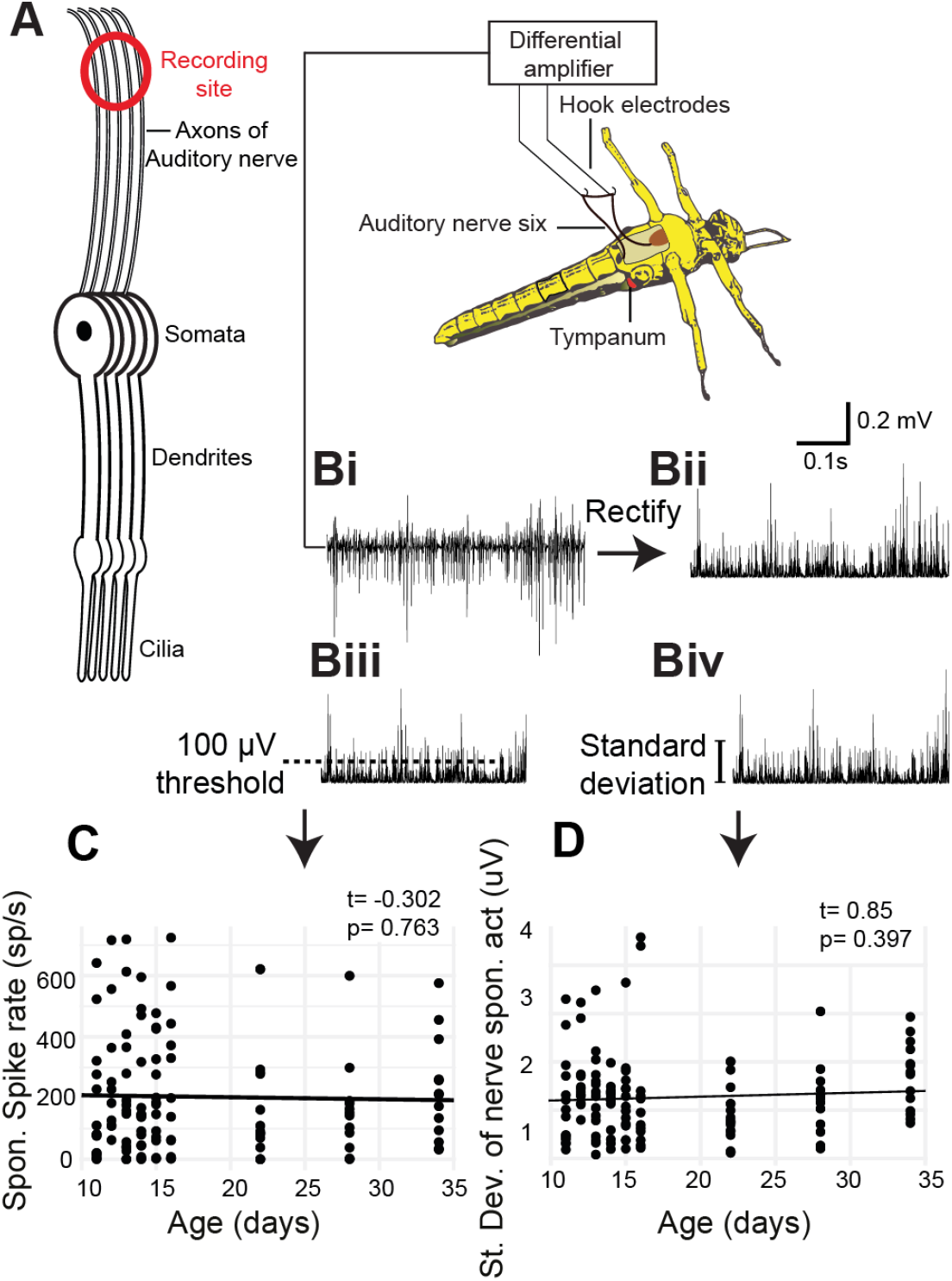
Metabolic activity of Müller’s organ measured as output from the auditory nerve. A Experimental setup of hook electrode recordings. **Bi** Spontaneous potentials from the auditory nerve, **Bii** rectified potentials from the auditory nerve, **Biii** 100 μV threshold used to count spontaneous spikes and **Biv** standard deviation of the rectified nerve potentials. **C**. Quantification of the number of spikes as a function of age. **D**. Quantification of the standard deviation of the rectified nerve potential as a function of age.

### Electrophysiological properties of auditory neurons

Auditory neurons of insects pass cations through their transduction channels, from the receptor lymph cavity, an enclosure of a high electrochemical gradient (analogous to the scala media endolymph in the cochlea). We pharmacologically blocked spiking activity of the auditory neurons to only observe the activity of the transduction channels (Figure 3A). The transduction channels of auditory neurons spontaneously open even at rest to produce discrete depolarisations (Figure 3Bi, Bii). Both the amount of current that passed through the transduction channels (Figure 3Ci) and the amplitude of the discrete depolarisations (Figure 3Cii) decreased as a function of age (LM: t_(200)_=−2.037, p=0.0429 (Cohen’s d (comparing day 10 with day 34)=0.837) and t_(201)_=−3.066, p=0.0025) (Cohen’s d (comparing day10 and day 34) = 0.758). The resting potential of the auditory neurons remained unchanged as a function of age (Figure 3Ciii (LM: t_(195)_=−0.044, p=0.965). Next we stimulated with 3 kHz tone across a range of sound amplitudes and recorded the transduction current whilst clamping the auditory neuron at −100 mV to increase the driving force of cations into the neuron (Figure 3D). We fitted four part log linear functions to the transduction current as a function of the sound amplitude for each recording (Figure 3E, Log Linear fits are only shown for the average data here). The inflection point was not different between young and old locusts’ auditory neurons (Figure 3Fi) (LM: t_(203)_=0.985, p=0.325) but the hill coefficient tended to change (Figure 3Fii) (LM: t_(203)_=1.696, p=0.091). The maximum transduction current, produced with the largest sound amplitude (110 dB SPL), allows a measure of the electrochemical potential of the receptor lymph cavity (Warren et al., 2021). The maximum transduction current did not significantly decrease as a function age (Figure 3Fiii) (LM: t_(230)_= −1.42, p=0.156.

**Figure 3.**
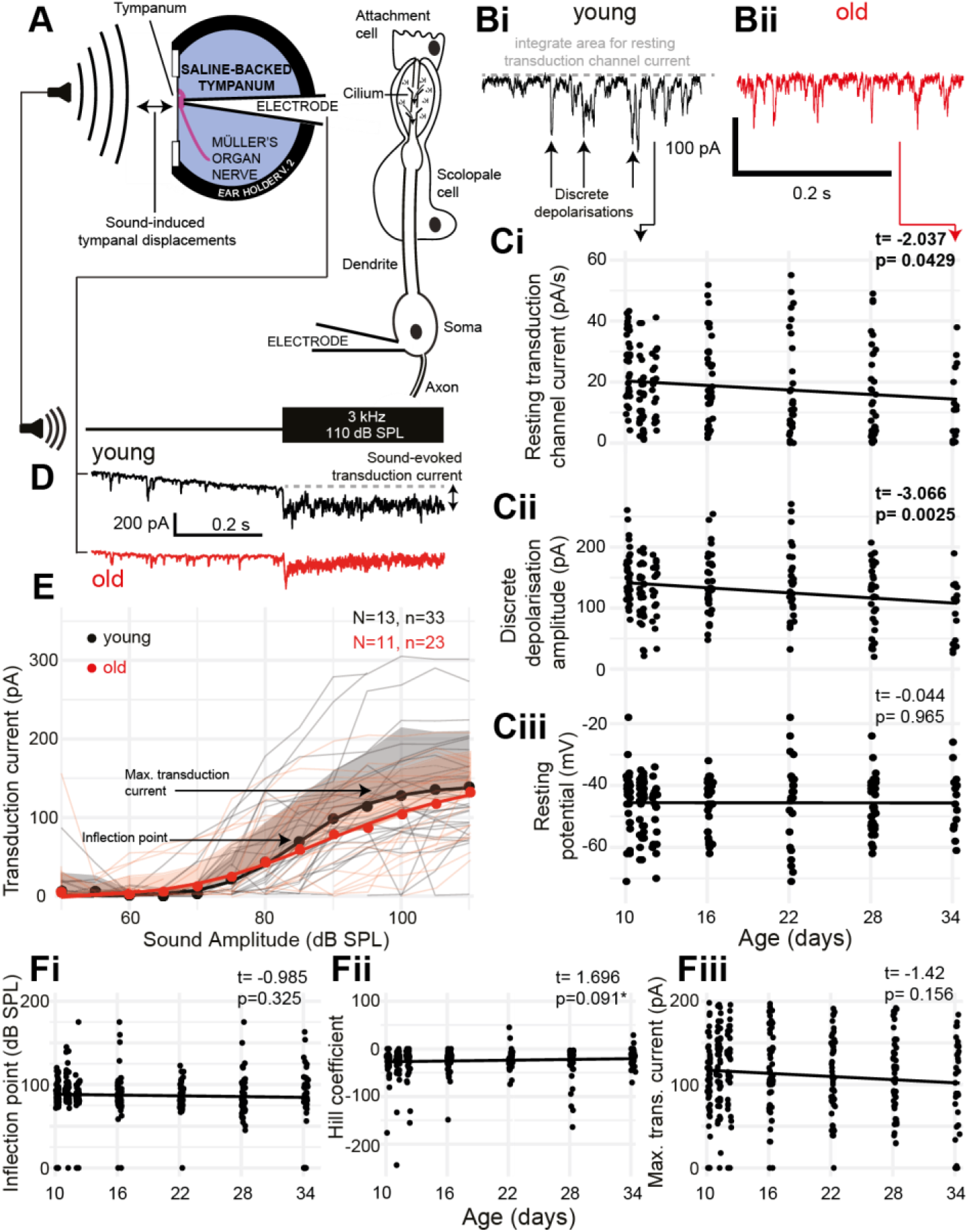
Properties of aging auditory neurons. **A**. Experimental setup and recording configuration. **B**. Voltage-clamp trace of auditory neuron resting transduction channel stochastic openings (discrete depolarisations) from **Bi** young (black) and **Bii** old (red) (34 days post final moult) locust ears. Ci. Quantification of the amount of resting current flowing through the transduction channels as a function of age. **Cii** Quantification of the amplitude of discrete depolarisations as a function of age. **Ciii**. Resting potential of the auditory neurons as a function of age. **D** Example of sound-evoked depolarisation measured in voltage-clamp mode for young (black) and old (red) locusts. **E**. Quantification of the transduction current as a function of sound amplitude for young (black) and old (red) locusts. Dots are average, solid lines are a log-linear four compartment fit to the data, shaded area is the positive standard deviation for control (grey) and aged (pink) locusts. Thin lines are the measurements from individual locusts for young (grey) and old (pink) locusts. **Fi** Inflection point (as shown in E) of the log-linear fits to transduction currents from individual locusts across ages. **Fii** Hill coefficient (shown in E) of the log-linear fits to transduction currents from individual locusts across ages **Fiii** Maximum transduction current (shown in E) of the log-linear fits to transduction currents from individual locusts across ages.

### Individual differences in metabolism and sound-evoked auditory nerve activity

If metabolism is causative for age-related auditory decline then auditory organs with a lower metabolism should have decreased auditory function regardless of age (and *visa versa*). We exploited the natural variation in metabolism and auditory nerve response to measure both in the same locust’s ear. Hook recordings were performed first to measure the auditory nerve response, then the same ear was extracted and its metabolism quantified using the resazurin metabolic assay (Fig. 4A) within 10 minutes of finishing the hook electrode recording (Figure 4B). We found no correlation between the auditory nerve response and the rate of metabolism (Figure 4C) (t_(39)_=0.267, p=0.79) even with a large variation in both measurements (R^2^=0.043).

**Figure 4.**
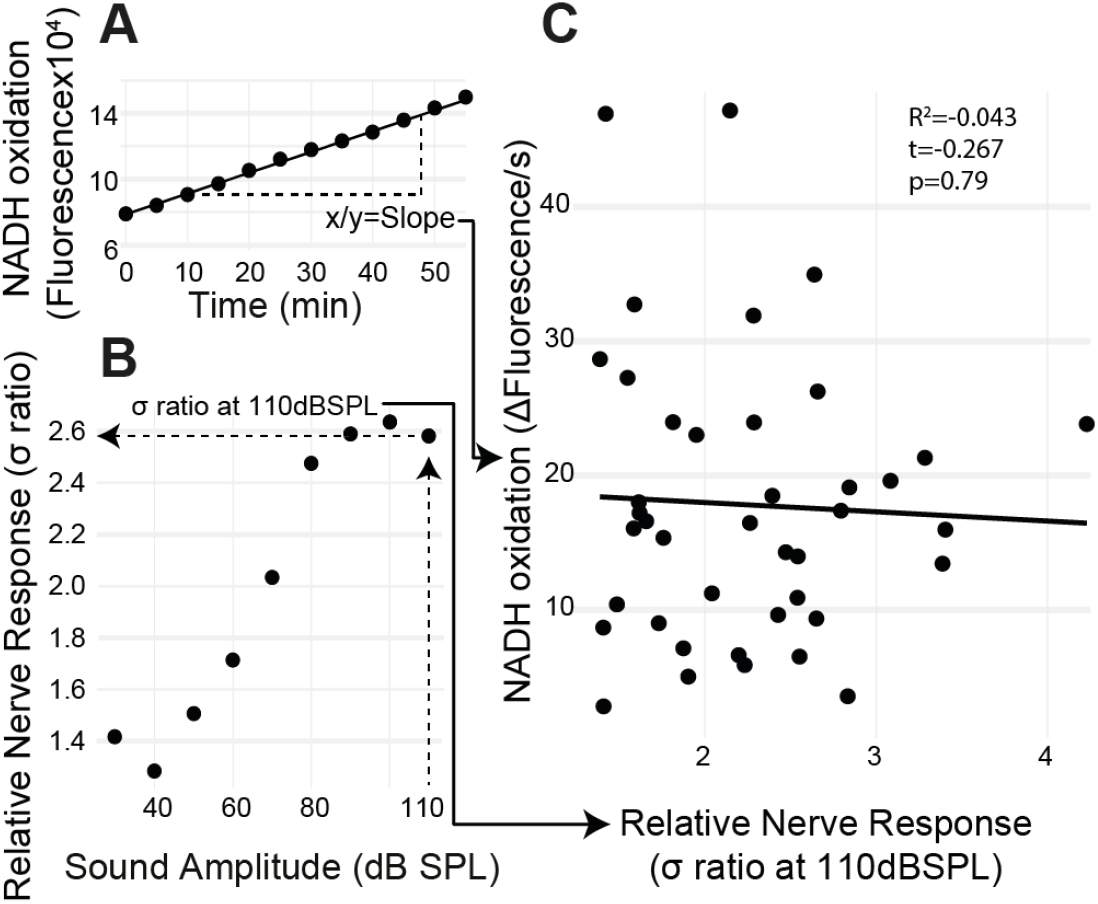
Ear-specific correlation of sound-evoked nerve activity and metabolism. **A**. Metabolic rate was measured by quantifying the slope of fluorescent changes as a function on time. **B**. The maximum nerve response σ ratio was measured at 110 dB SPL. **C**. Both measurements were plotted for each ear with no correlation between auditory nerve function and metabolism.

### Cold-rearing locusts increased Müller’s organ metabolism

We exploited the ectothermic nature of the locust to manipulate chemical reaction rates through altering the environmental temperature. We raised locusts either in standard environmental conditions (32:25°C day:night) or cold conditions (12:6°C day:night). We successfully altered the metabolic rate of both muscle tissue in foreleg tibia (Fig. 5A, t_(27)_=5.89, p=2×10^−7^; Cohen’s d= −2.08) and Müller’s organ, (Figure 5B, t_(43)_=3.67 p=0.0004; Cohen’s d=−0.57). The sound-evoked response of the auditory nerves were different between locusts raised in cold conditions (Figure 5C, F Statistic=6.12). This difference was manifested by a lower maximum asymptote of the auditory nerve response in cold-reared locusts, opposite to the increase of metabolic rate in Müller’s organs (t_(4)_=−3.37, p=0.00086, Cohen’s d=0.793 (model fit at 110 dB SPL)). The other three parameters of the four-part log-linear fit were not different (Hill coefficient t_(4)_=0.22, p= 0.82; Lower asymptote t_(4)_=0.20, p=0.84; inflection point t_(4)_=−1.74, p=0.08).

**Figure 5.**
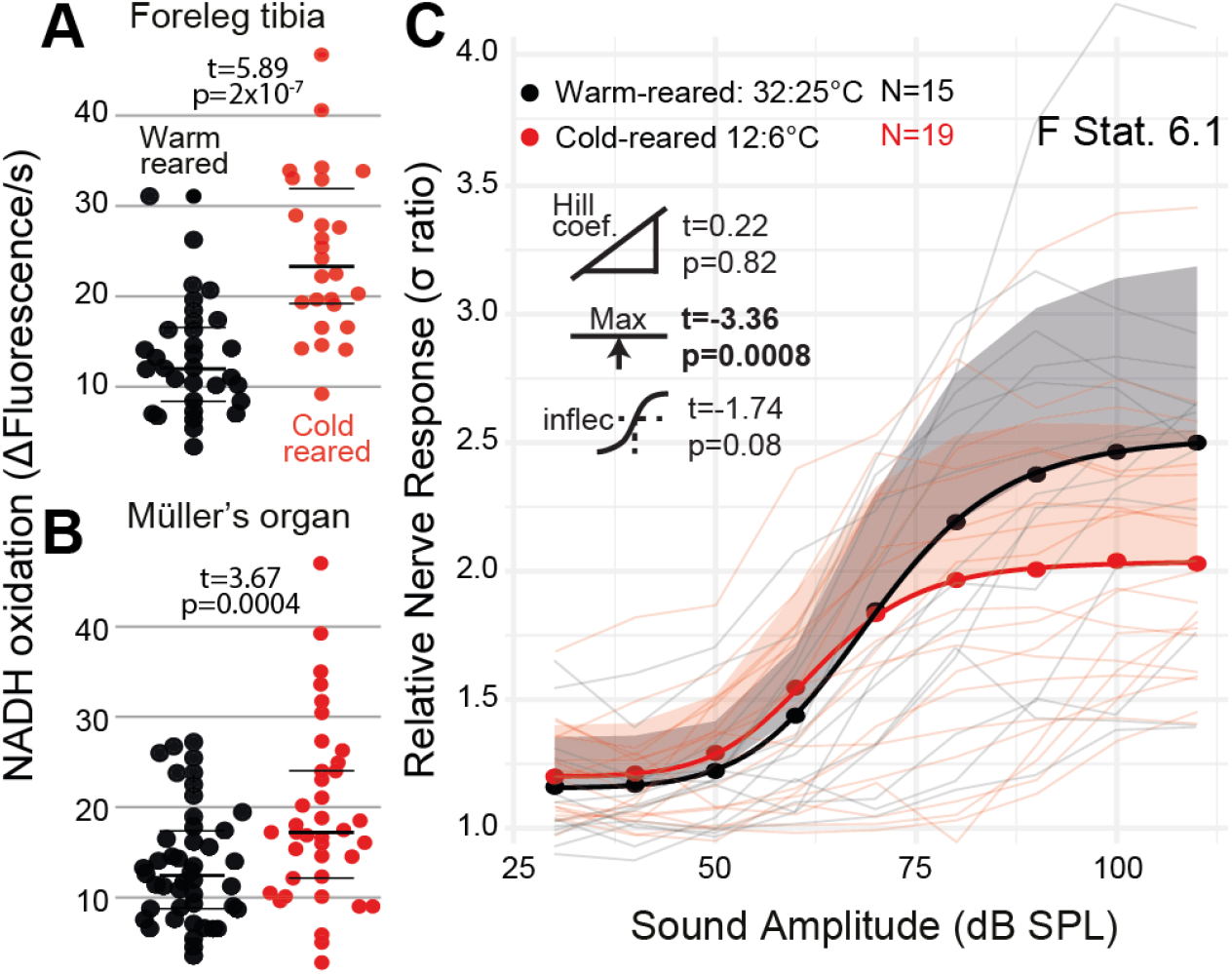
Quantification of metabolic rate and auditory nerve function in cold-reared locusts. **A**. The metabolic rate of reduction of resazurin through NADH oxidation increased in foreleg tibia in normal (32:25 °C day:night) conditions and cold (12:6 °C day:night) conditions. **B**. The metabolic rate of Müller’s organs in warm- and cold-reared locusts. **C**. Sound-evoked responses of locusts reared in warm- and cold conditions with respect to dB SPL. Responses from individual locusts are plotted as thin lines (pink=cold-reared, black=warm-reared), averages for warm- and cold-reared locusts are dots. The positive standard deviation of the averages are shaded regions and the solid thick lines are the four-part log-linear fit to the individual responses.

### Starving locusts reduced Müller’s organ metabolism

We starved locusts for five days which failed to reduce the metabolic rate of muscle tissue in foreleg tibia but the metabolic rate was reduced Müller’s organs compared to locusts fed *ab libitum* (Fig. 6A t_(27)_=0.22, p=0.824; 6B t_28_=−3.59, p=0.001; Cohen’s d= 1.31). The sound-evoked response of the auditory nerves across sound pressure levels was not significantly different between starved and fed locusts (Figure 6C, F statistic 0.17) for any of the four parameters of the log linear fit (Maximum asymptote t_(4)_=0.29, p=0.77; Hill coefficient t_(4)_=0.59, p=0.56; Lower asymptote t_(4)_= 0.01, p= 0.99; inflection point t_(4)_=0.12, p=0.90).

**Figure 6.**
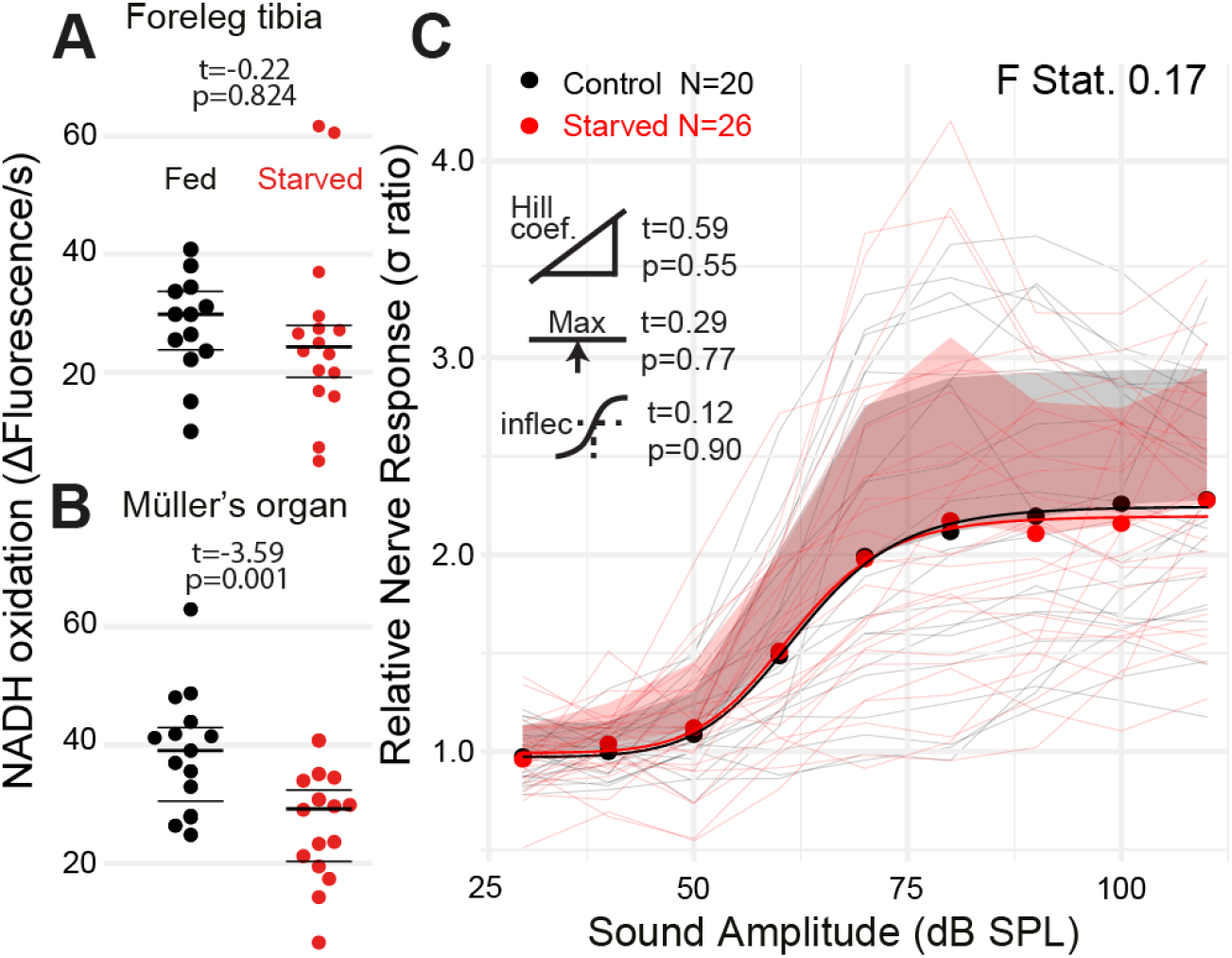
Quantification of metabolic rate and auditory nerve function in starved locusts. **A**. The metabolic rate of reduction of resazurin through NADH oxidation increased in foreleg tibia in locusts fed *ab libitum* and starved locusts **B**. The metabolic rate of Müller’s organs in locusts fed *ab libitum* and starved locusts. **C**. Sound-evoked responses of locusts fed *ab libitum* and starved locusts. Responses from individual locusts are plotted as thin lines (pink=starved, black=fed), averages for starved and fed locusts are dots. The positive standard deviation of the averages are shaded regions and the solid thick lines are the four-part log-linear fit to the individual responses.

### RNA expression changes with age

To further examine the role of metabolism in age associated auditory decline, we compared the transcriptome of the Locust’s Müller’s organs between young and old locusts. Transcriptomic results showed 1109 differentially expressed genes between young and old locusts. 486 genes were upregulated in the younger locusts, whilst 623 genes were upregulated in the older locusts. After annotating the *Schistocerca gregaria* genome with GO Terms, we performed GO enrichment on the differentially expressed genes. We obtained 16 GO terms for cellular component, 20 for molecular function, and 85 for biological process. The top cellular component GO terms were associated with the extracellular space.

Many of the most enriched biological GO terms were for terms associated with chitin development, such as cuticle development, chitin-based cuticle development, and chitin metabolic process. This is due to the large number of differentially expressed genes associated with chitin based process. For example, amongst the top 20 most significant genes were cuticle protein 65 (LOC126291912 & LOC126355498), cuticle protein 16.5 (LOC126291616) and cuticle protein 5 (LOC126355435).

Although immune system process were not enriched in this analysis, a number of GO terms associated with immunity were found to be significantly enriched, including defense response to gram-positive bacterium, response to external biotic stimulus and activation of innate immune response (Figure 7). These genes associated with these GO terms are generally upregulated in the old locusts. In addition, amongst the top 100 differentially expressed genes were genes including hemocytin (LOC126356178) two mucin-19 genes (LOC126295440 & LOC126272492). Additionally to mucin-19, we find mucin-5AC (*MUC5ac*)(LOC126281354) differentially expressed. *MUC5ac* overexpression has been associated with hearing loss (Lee *et al*. 2015), although paradoxiacally is upregulated in the young locust. We also found enrichment for the inflammatory response GO term, with genes such as Mucin-17 (LOC126337069), nuclear factor NF-kappa=B pp1010 subunit (LOC126282030) and peptidoglycan recognition protein 1 (LOC126267215 & LOC126267215).

**Figure 7.**
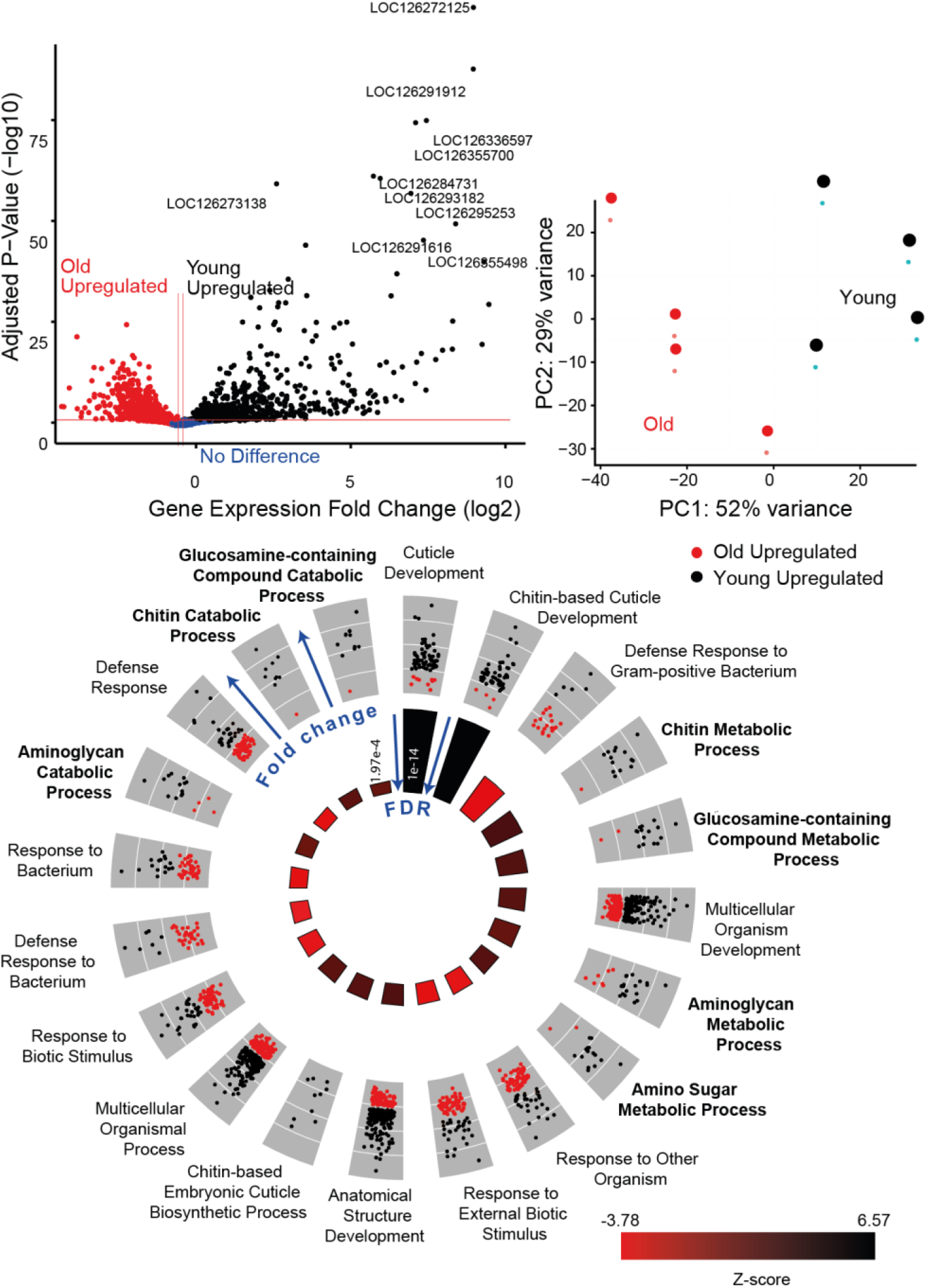
Transcriptomic results for young (black) versus old locusts (red) Müller’s organs. **A**. Volcano plot comparing log fold p-value and log fold change, annotated with the top 10 differentially expressed gene’s LOC numbers **B**. PCA of gene expression counts. **C**. GO circle plot of top 20 enriched biological process GO terms. Each gene’s associated with the GO term is plotted in relation to its fold change, either being upregulated in old (red) or upregulated in young (black). In the inner circle, plots for the z-score are reported, which give an indication as to whether the group of genes associated with a particular GO term are more upregulated in young locust ears (black) or old (red), with the size of the bar indicating the size of the FDR (larger bars being more significant).

Other interesting enriched biological process GO terms included cytolysis and response to estrogen. Additionally, we identified some genes which had previously been implicated in hearing loss. Some of these genes include *RXR* (LOC126272034), and *eEF2* (LOC126272205), *SLC52A3* (LOC126272245).

Focusing primarily on metabolism, we found enriched GO terms associated with metabolic processes including aminoglycan metabolic process, amino sugar catabolic process and carbohydrate derivative catabolic process (Figure 7). In total, 274 metabolic process associated genes were found to be differentially expressed. We found some NAD metabolism associated genes to be differentially expressed; L-lactate dehydrogenase (*LDH*) (LOC126266619) and pyruvate kinase (LOC126274964) which both show increased expression with age, and glycerol-3-phosphate dehydrogenase (LOC126266753) which shows decreased expression with age.

## Discussion

Metabolism is hypothesised to be responsible for age-related hearing loss. However, metabolic rate has never been measured as a function of age in an auditory organ and neither has metabolism been manipulated to establish a causative role. Using an insect model, we measured metabolic rate in an auditory organ through quantifying the reduction of resazurin, a redox signaller and key coenzyme for glycolysis and oxidative phosphorylation. We then manipulated metabolic rate in an attempt to find a causative role of metabolism in age-related hearing loss. Finally, we measured gene expression changes as a function of age in Müller’s organ.

We found that metabolic rate is lower in Müller’s organs in older locusts paralleling that found across whole-body basal metabolism in a range of animals (*Drosophila*: Fiorino et al., 2018; Guppies: Imai et al., 2022; Mice: Haramizu et al., 2011; Humans Henry, 2000). The slowing of metabolism later in a lifespan is also paralleled by basal metabolic measurements in humans (Henry, 2000). Whilst it is not surprising that metabolism decreased in Müller’s organs with age it could also be hypothesised that auditory tissue evolved a special resilience against metabolic-dependant age-associated deterioration due to the selective advantage of sensitive hearing for predator avoidance and mate finding. We predicted that interfering with metabolically demanding processes of the auditory neurons, through application of voltage-gated ion channel blockers and chordotonal organ-specific insecticide, pymetrozine, would result in a decrease in metabolism. However, like the mammalian cochlea (Waldhaus et al., 2015), the primary auditory receptors only make up 5% of cells in the auditory organ (~80 out of 1500 (Blockley et al., 2022)). It appears that the auditory receptors combined metabolic demands of the primary sensory receptors do not outweigh those of the supporting cells.

Although the number of auditory neurons decreases as a function of age (Blockley et al., 2022) the width of the auditory nerve and the number of Schwann cells engulfing it does not change with age (Blockley et al., 2022). Here we found that there is no age-dependence in the number of spontaneous spikes travelling along the auditory nerve. At the level of the auditory receptors we found a decrease in passing current through the transduction channels which mirrors that found in mouse hair cells (Jeng et al., 2021). We believe this could be a common mechanism to maintain the electrochemical potential of the receptor lymph to compensate for an age-dependant decrease (albeit mild in this this study t=−1.42) in the performance of ion-pumping scolopale cells or, in the case of mammals, cells of the lateral wall in the cochlea. The resting membrane potential of the auditory neurons is well-maintained into old age, similar to mouse hair cells (Jeng et al., 2021). Altogether, the metabolic demands of the auditory neurons appear to remain fairly constant over its lifespan, with a possible mechanism to compensate for age-related decline in the performance of ion-pumping cells.

Here, we wished to determine if whole auditory organ metabolism directly determined its performance or if metabolic performance is simply a correlative proxy for the age. Decreasing metabolism through dietary restriction is shown to extend lifespan in diverse organisms (McCay et al., 1935; Anderson et al., 2003; Klass, 1977; Partridge et al., 1987) presumably due to a decrease of oxidative stress, enhanced mitochondrial efficiency (Ruetenik & Barrientos, 2015), decreased DNA damage (Haley-Zitlin & Richardson, 1993) and a decreased immune response (Wu et al., 2019). At the same time a more metabolically active auditory organ would be expected to perform better, especially if this equates to an increased electrochemical gradient across the transduction channels (Schmiedt et al., 2002). Here, we found no correlation between the metabolic state of single Müller’s organs and its sound-evoked potentials. We also found no concurrent changes with auditory performance of Müller’s organ and its metabolism when we altered metabolic rate of Müller’s organ through dietary restriction. If anything, we found that whilst metabolism increased in in the cold reared Locusts, the maximum auditory nerve activity conversely decreased.

We believe our findings can best be explained by an age-correlated, but not causative role of metabolism in hearing loss. A useful example are the sex-differences in human hearing and metabolism. The metabolic rate of human males is higher than females (Shapiro et al., 1980; Gagnon & Kenny, 2012). If we assume that these metabolic sex differences extend to the cochlea we would expect that males have better hearing than females. In fact, not only are auditory thresholds similar in adolescence females - female hearing actually grows to outperform that of males (Park et al., 2016). This could be explained – albeit not exclusively - by higher oxidative stress in males that leads to accumulating damage, and performance, of the auditory system (Ide et al., 2002). By comparison, our findings of the co-occurrence of a decline in auditory function and metabolism could be explained by cumulative damage caused by oxidative stress and free radical production. The actual metabolic output of Müller’s organ does not determine its auditory function.

Whilst our transcriptomic results showed enrichment for some metabolic processes such as aminoglycan metabolic process, amino sugar catabolic process and carbohydrate derivative catabolic process, we also found enrichment for a number of other ageing processes. One aging hallmark - chronic inflammation - is contributed to by oxidative stress, age related changes in the inflammatory cytokine network and cellular senescence (Kim & Song, 2014). Immune responses genes were clearly upregulated in older Müller’s organs paralleling that found in the mouse cochlea (Schubert et al., 2022). We also found genes enriched for cytolysis to be differentially expressed between young and old locust auditory organs. These findings add further evidence for the multi-process nature of age related auditory decline.

We found a number of differentially expressed genes which have been implicated with hearing loss in mammalian systems. For example, *RXR*, a nuclear receptor whose expression declines with age, have been touted as potential therapeutic targets for the inner ear (Kwak *et al*. 2019). Here we found *RXR*’s expression is reduced in older locust Muller’s organs compared to young. In mammals the autophagy gene *eEF2*’s repression can alleviate many ageing phenotypes, and is found to be upregulated in severe hearing loss (Peng *et al*. 2022). Between young and old locusts we found *eEF2* to be upregulated in the older locusts. A final example is *LDH*. *LDH* reversibly catalyses the reaction of lactate to pyruvate. The *LDH* gene upregulated in aged Locusts is most similar to *LDH-A*, as was also found in mice (Ross *et al*. 2010). This increases the pyruvate to lactate reaction, which has previously been implicated in human age related hearing loss (Tian *et al*. 2020). The enriched GO term for Melanin biosynthesis also appears to increase with age, which may have a protective quality - melanin precursors have been shown to prevent both age-related and noise-induced hearing loss in mice (Murillo-Cuesta et al., 2009). In humans, black individuals also have a lower risk of hearing loss than their white counterparts, perhaps due to having increased inner ear melanin (Agrawal et al., 2008).

Mammalian studies have shown that different cell types have differential expression of aging-related genes, some of which may contribute to auditory decline (Schubert et al., 2022). In the Müller’s Organ, there are thought to be at least 4 different cell types. To help further elucidate the contribution of each of these cell types, a more targeted characterisation is needed for each of these cells. The inclusion of a large number of chitin genes shows the difficulty in performing whole tissue RNA sequencing. By performing RNAseq on the whole Müller’s Organ and a portion of attached tympanum, chitinous tissue contributes largely to the overall differential expression. Many of the most enriched biological GO terms were involved in chitin development – this is to be expected as chitin production decreases significantly as a function of age post final-moult (Candy & Kilby, 1962). Single cell sequencing may provide an opportunity to characterise the distinct cell types of the auditory organ, and elucidate how they each individually contribute to age related hearing loss. This will be particularly useful for understanding changes in expression within Scolopale cells and sensory neurons, as currently their metabolic output (and potentially changes in expression of related genes) is potentially being dwarfed by supporting cells.

## Conclusion

We measured a decrease in metabolism in the auditory organ of the desert locust as a function lifespan. Auditory neurons showed little age-related changes in function and no dominating contribution to metabolic rate. We found no correlation between the metabolic output of individual Müller’s organs and their ability to transduce sound-evoked potentials. We altered metabolic rate of the auditory Müller’s organ through dietary restriction and cold-rearing and found no change or opposite changes in the magnitude sound-evoked response of Müller’s organ. Age-related changes in gene expression suggest that other processes, outside of metabolism, dominate the age-related decline of Müller’s organ.

## Methods

### Locust husbandry and conditioning

Desert locusts, *Schistocerca gregaria* (mixed sex) were reared in crowded gregarious conditions 150-250 in 60 cm^3^ cages in their fast-aging gregarious state, where they can live up to two months. They had a 12-h light/dark cycle at 32:25°C and were fed on a combination of fresh wheat and bran *ad libitum*. The gregarious state of the desert locust contrasts with their isolated solitarious state where they live for up to nine months. The founding progeny of the Leicester Labs strain were solitary copulating adults collected at Akjoujt station ~250 km North East from Nouakchott, Mauritania in May 2015.

After hatching desert locusts go through five moults before they are adults with functional wings. For the longitudinal metabolic assay (Figure 1) ~20 locusts (mixed sex) were taken within 24 hours of their final moult from their larger cages (detailed above) and placed in plastic aquarium tubs (40×20×30 cm) with *ab libitum* wheat and milled bran.

For longitudinal hook electrode and patch clamp recordings (Figures 2 and 3) locusts (mixed sex) were taken 10 days post their last moult and had their wings clipped at their base. These locusts were when aged in aquarium tubs as described above.

The locusts to test the effect of TTX&TEA, electron transport chain blockers and pymetrozine (Figure 1Bi, ii, iii) were all 10 days post their last moult. The locusts used to test causality of metabolism on hearing loss (Figures 4) were all 10 days post their imaginal moult and male. Locusts were 10 days post their last moult before entering the conditioning pipeline for rearing in cold conditions for 15 days at 7:15°C or starving 9water only) for five days.

For Fig. 5, Locusts 10 days past final moult were reared in a 12hr light/dark cycle 12:6°C, for 14 days. Control locusts were reared under standard conditions as previously described for the same duration. Auditory nerve recordings and ear extractions were carried out at room temperature. For Fig. 6, male locusts, 10 days past their final moult, were taken and raised under normal conditions, but without food for 7 days. Control animals were raised under normal conditions, fed *ad libitum*.

### Metabolic assay

A 0.02% (weight/volume) solution of resazurin sodium salt was dissolved in locust saline. Locust saline had the following concentrations (in mM): 185 NaCl, 10 KCl, 2 MgCl2, 2 CaCl2, 10 HEPES, 10 Trehalose, 10 Glucose. The saline was adjusted to pH 7.2 using NaOH. Three other solutions were used as positive control, or to test the metabolic involvement of the auditory neurons. For the positive control we used three electron transport chain blockers dissolved in resazurin sodium saline solution: 1 mM sodium azide, 0.5 μM antimycin, 0.5 μm rotenone. To selectively target the spiking activity of auditory neurons we used 90 nM Tetrodotoxin (TTX) and 20 mM Tetraethylammonium (TEA) and to silence auditory neurons completely (see Warren and Matheson, 2018) we used 30 μM pymetrozine all dissolved in resazurin sodium saline solution.

Volumes of 30 μl of the resazurin containing saline were pipetted into a 384 cell culture microplate with flat bottom and transparent lid (781086, Greinerbio-one) on ice. Müller’s organs of locusts were extracted by inserting Dumont #5 forceps (501985, World Precision Instruments) so that each forcep point penetrated the tympanum (by about 0.5 mm) flanking the folded body. The forceps clasped Müller’s organ before pulling it away from the tympanum. The forcep was inspected for confirm extraction of Müller’s organ before repeating the process for the other ear of the locust. Both Müller’s organs were placed into a single well for Figure 1A. We realised that this assay was sensitive for single ears and for Figures 1Bi, ii, iii single Müller’s organs were placed in each well. To measure the metabolic rate of tissue separate from Müller’s organ we took the fore leg tibia and cut it on the tibia side of its joint. We placed it distal side up in the 384 Microwell plate well.

Resazurin (a weakly fluorescent blue dye) is oxidized by NADH into highly fluorescent resorufin (excitation/emission 571/584 nm). Using the FLUOstar Omega BMG Microplate reader we excited each well using an excitation filter at 544 nm and measured through an emission filter of 590 nm. Fluorescent intensity measurements with a gain of 1800 were taken every 5 minutes for 12 cycles. The increase in fluorescence were fitted with a regression line in Mars analysis software (BMG Labtech) to give an increase in fluorescent units per second.

### In vivo hook electrode recordings from auditory nerve six

Locusts were secured ventral side up with their thorax wedged in a plasticine channel and their legs splayed and held down with plasticine. A section of the second and third ventral thoracic segment was cut with a fine razor blade and removed with fine forceps. Tracheal air sacks were removed to expose nerve six and the metathoracic ganglia. This preparation left the abdomen, including the 1^st^ segment where the ears reside intact. Thus, maintaining the operation of the ear *in vivo*. Hook electrodes constructed from silver wire 18 μm diameter (AG549311, Advent Research Materials Ltd) were hooked under the nerve and the nerve was lifted out of the haemolymph. For Figure 2 a mixture of 70% Vaseline and 30% paraffin oil was applied through a syringe to coat the auditory nerve to stop it drying out. For figures 4, 5, 6 the nerve was not coated with a mixture of vasaline and paraffin oil. Locust mounting and recordings took ~ 15 minutes for each locust. For figure 4 Müller’s organs were extracted as detailed above directly after hook electrode recordings were completed (within 10 minutes). Signals were amplified 1,000 times by a differential amplifier (Neurolog System) then filtered with a 500 Hz high pass filter and a 50 kHz low pass filter. This amplified and filtered data was sampled at 25 kHz by Spike2 (version 8) software running on Windows (version 10). To quantify the compound spiking activity of the auditory nerve we used Matlab (Version R2020a, Mathworks Inc.) and rectified the nerve signal and integrated the area underneath. We computed this for the 0.5 s of sound-evoked neural activity and for 60 s background nerve activity before the tones and the background activity between the tones. To compute the σ ratio we rectified the nerve signal and removed any DC offset. We then divided the sound-evoked response by the background neural activity. For the Starved locusts (Fig.6A), the locust treatment was blinded to the experimenter until all data was collected and analysed. Blinding was not possible for the Cold reared experiments (Fig.5A), as there was an observable phenotypic difference in cuticle colour and texture between the animals raised at different temperatures.

### Dissection of Müller’s Organ and isolation of Group-III auditory neurons

Whole cell patch clamp recordings were performed on group-III auditory neurons because they form the majority of auditory neurons of Müller’s organ (~46 out of ~80) (Fig. 2A) (Jacobs et al., 1999), they are the most sensitive auditory neurons of Müller’s organ (Römer, 1976) and are broadly tuned to the 3 kHz we used for noise-exposure (Warren and Matheson, 2018). For intracellular patch-clamp recordings from individual auditory neurons the abdominal ear, including Müller’s Organ attached to the internal side of the tympanum, was excised from the first abdominal segment, by cutting around the small rim of cuticle surrounding the tympanum with a fine razor blade. Trachea and the auditory nerve (Nerve 6) were cut with fine scissors (5200-00, Fine Science Tools), and the trachea and connective tissue removed with fine forceps. This preparation allowed perfusion of saline to the internal side of the tympanum, necessary for water-immersion optics for visualizing Müller’s Organ and the auditory neurons to be patch-clamped, and concurrent acoustic stimulation to the dry external side of the tympanum. The inside of the tympanum including Müller’s Organ was constantly perfused in extracellular saline. Dissection, protease and recordings took ~ 60 minutes for each locust ear.

To expose Group-III auditory neurons for patch-clamp recordings, a solution of collagenase (0.5 mg/ml) and hyaluronidase (0.5 mg/ml) (C5138, H2126, Sigma Aldrich) in extracellular saline was applied onto the medial-dorsal border of Müller’s Organ through a wide (12 μm) patch pipette to digest the capsule enclosing Müller’s Organ and the Schwann cells surrounding the auditory neurons. Gentle suction was used through the same pipette to remove the softened material and expose the membrane of Group-III auditory neurons. The somata were visualized with a Cerna mini microscope (SFM2, Thor Labs), equipped with infrared LED light source and a water immersion objective (NIR Apo, 40x, 0.8 numerical aperture, 3.5 mm working distance, Nikon) and multiple other custom modifications. For a full breakdown of the microscope components and how to construct a custom patch-clamp microscope for ~£12ksee: https://le.ac.uk/people/benjamin-warren.

### Whole-cell patch-clamp recordings

Electrodes with tip resistances between 3 and 4 MΩ were fashioned from borosilicate class (0.86 mm inner diameter, 1.5 mm outer diameter; GB150-8P, Science Products GmbH) with a vertical pipette puller (PC-100, Narishige). Recording pipettes were filled with intracellular saline containing the following (in mM): 170 K-aspartate, 4 NaCl, 2 MgCl2, 1 CaCl2, 10 HEPES, 10 EGTA. 20 TEACl. Intracellular tetraethylammonium chloride (TEA) was used to block K^+^ channels necessary for isolation the transduction. To further isolate and increase the transduction current we also blocked voltage-gated sodium channels with 90 nM Tetrodotoxin (TTX) in the extracellular saline. The addition of ATP to the intracellular saline did not alter the electrophysiology of the recordings so was omitted. During experiments, Müller’s Organs were perfused constantly with locust saline (same as that used for metabolic assays). The saline was adjusted to pH 7.2 using NaOH. The osmolality of the intracellular and extracellular salines’ were 417 and 432 mOsm, respectively.

Whole-cell voltage-clamp recordings were performed with an EPC10-USB patch-clamp amplifier (HEKA-Elektronik) controlled by the program Patchmaster (version 2x90.2, HEKA-Elektronik) running under Microsoft Windows (version 7). Electrophysiological data were sampled at 50 kHz. Voltage-clamp recordings were low-pass filtered at 2.9 kHz with a four-pole Bessel filter. Compensation of the offset potential were performed using the “automatic mode” of the EPC10 amplifier and the capacitive current was compensated manually. The calculated liquid junction potential between the intracellular and extracellular solutions was also compensated (15.6 mV; calculated with Patcher’s-PowerTools plug-in from www3.mpibpc.mpg.de/groups/neher/index.php?page=software). Series resistance was compensated at 77% with a time constant of 100 μs. The resting potential was measured directly (within 10s) after whole-cell recordings were established by changing the clamped voltage until the current was zero. We measured the amplitude of the discrete depolarisations by measuring the largest three discrete depolarisations at a −100 mV holding potential. To measure the standing current we measured spontaneous currents for 0.5 s at a holding potential of −60 mV. We integrated the area underneath the current at baseline (i.e. periods with no discrete depolarisations, See Figure 3Ci). For all patch-clamp electrophysiological analysis we used Igor Pro 9, (Wavemetrics Inc.).

### RNA extraction

Müller’s organs from the tympanal ears of locusts were extracted as described above in *metabolic assay*. Four groups of 15 male locusts ~10 days post their last moult and four groups of 15 male locusts ~24 days post their last moult were used (8 groups in total). Müller’s organs were snap frozen by wiping them onto a frozen pestle that rested in an Eppendorf tube submerged in liquid nitrogen. The frozen samples were then homogenised by hand with the pestle for 3 minutes. 10 μL of Trizol was then added to the samples, and homogenised for a further 3 minutes at room temperature. Next 490ul of Trizol was added to the samples and mixed gently by pipetting. Samples were then centrifuged for 10 minutes at 12,000g. Supernatant of the sample was collected, and left to incubate at room temperature for 5 minutes. 100 μl of chloroform was added to the samples, and the sample was then vortexed. Samples were incubated for 3 minutes, and then centrifuged for a further 20 minutes at 12,000g. The aqueous phase was then separated into a fresh RNase-free tube, along with 1 μl of glycogen and 250 μl of isopropanol. The samples were then mixed gently and left to incubate at room temperature. Samples were then centrifuged for 10 minutes at 12,000g. The supernatant was then removed and the RNA was then washed with 500ul of 75% ethanol. The pellet was then dislodged via vortexing, before centrifuging again at 7,500g for 5 minutes. Supernatant was again removed and the pellet was air dried until it began to turn clear (approx. 2 minutes). Samples were then suspended in 10 μl of RNase-free water and incubated for 5 minutes at 55 °C. RNA was quantified via nanodrop and quality control was carried out on bioanalyser. Samples were then stored at −80°C. Samples were treated with DNAse as described in Invitrogen™ RNAqueous™-Micro Total RNA Isolation Kit, if required.

### Bioinformatic analysis

Samples were sequenced and quality trimmed on the Illumina platform by Novogene sequencing (Cambridge) with 150 bp paired ends at a read depth of 60 million. Read quality was checked using FastQC (v0.11.5) (Andrews & Simon 2015). RNA reads were aligned to the iqSchGreg1.2 reference genome (NCBI) using STAR 2-pass method (v2.7.9a) (Dobin et al. 2013), and the resultant bam files were sorted using Samtools (v1.9) (Li et al. 2009). Gene counts were generated using HTSeq (Anders et al. 2015), and differential expression analysis was performed using DESeq2 (v1.28.1) (Love et al. 2014) on Rstudio (4.2.1), with p-adj < 0.05. GO terms were generated for genes in the SchGreg1.2 using eggNOG 5.0 web tool (Huerta Cepas et al. 2019). GO term enrichment was performed using topGO (Alexa & Rahnenfuhrer 2010) FDR < 0.05. GO term circle plots were generated using the GOPlot R package (Walter *et al*. 2015).

### Power analysis

We designed experiments that formed figures 2, 3, to have a power above 95%, which gives an ability to detect a difference between young and old or control and conditioned locusts or control and experimental Müller’s organs of 95% (if these series of experiments were run an infinite amount of times). Our false negative rate, or type II error probability is < 5% (1-power) (probability of not finding a difference that is there). Our false positive rate or type II error (probability of finding a difference that is not there) is determined by our p-values which was set at 0.05. In order to calculate the power, we used the raw data and effect size reported in Warren et al. (2020) for hook electrode recordings of spontaneous nerve activity (Figure 2). This was also carried out for whole-cell patch-clamp recordings from individual auditory neurons of Müller’s organ (Figure 3). There exists no analytical methodology for conducting power calculations on Linear Mixed Effect Models (LMEM). Therefore we generated a dataset simulated from the raw data of Warren et al. (2020), fitted a Linear Mixed Effect Model, then ran repeated simulations of the LMEM 1000 times. We used the proportion of times that the LMEM reported a difference to calculate the power. For this paper the experiments that measured differences in hook electrode spontaneous nerve activity and the spontaneous transduction channel openings (termed discrete depoarlisations) all had at least 95% power when using n numbers of 9 and 12 respectively. This simulated power analysis also, only applies for the effect size reported in Warren et al (2020) and may be lower if the actual effect size, in this paper, is reduced. Models were fitted in R (Version 2.4.3), on a Windows PC running Windows 10 using the package *LME4* (Bates et al., 2015) and simulations were run with the package *simr* (Green and Macleod, 2016).

As metabolism has never been measured in auditory tissue we had no effect size to base a power analysis for the collection of age-related changes in the metabolic rate of Müller’s organ. Instead we decided to collect ears from a high number of locust ears (~500) to stand the best chance of detecting a difference (Figure 1). We display Cohen’s d effect size in all data sets where a difference was detected so that this effect size can be used for future power calculations. This same approach was used for our attempt to find a correlation between sound-evoked auditory nerve activity and metabolism, exploiting the natural variation in individual Müller’s organs (Figure 4). To calculate the number of locusts required to detect differences in Müller’s organ function for locusts

### Statistical analysis

Throughout the manuscript n refers to the number of recorded neurons and N refers to the number of Müller’s Organ preparations used to achieve these recordings (i.e. n=10, N=6 means that 10 neurons were recorded from 6 Müller’s Organs). All n numbers are displayed on the figures for clarity. The Spread of the data is indicated by 1 standard deviation as the standard deviation indicates the spread of the data, unlike standard error. Median and Q1 and Q3 are displayed by bars when individual measurements are plotted. For all hook electrode recordings, 80% of patch-clamp recordings the treatment of the locust (noise-exposed or control) was blinded to the experimenter; lone working conditions, due to Covid restrictions, made complete blinding impossible. All data either remained blinded or was recoded to be completely blind when analysing the data to avoid unconscious bias.

To test for differences and interactions between control, noise-exposed and aged locusts we used either a linear model (LM) or Linear Mixed Effects Model (LMEM), with treatment and age as fixed effects, and Locust identity and SPL as a random intercept, when repeated measurements are reported. Models were fitted in R (Version 3.4.3) with the package *LME4* (Bates et al., 2015). The test statistic for these analyses (t) are reported with the degrees of freedom (in subscript) and p value, which are approximated using Satterthwaite equation (lmerTest package) (Kuznetsova et al., 2017). We report Cohen’s d effect size for significant differences. Curves were fitted to the data using the *drm* package in R for patch-clamp and hook electrode recordings (Ritz and Strebig, 2015). The *drm* package was also used to compute t and p values when comparing control and noise-exposed four part Log-Linear models. F statistics of the Log-Linear model fits were computed by excluding treatment (noise-exposed or control) as a factor. Higher F statistics donate a stronger effect of treatment.

In order to compare responses between control locusts and aged, starved or cold-reared locusts across SPLs we adopted an approach first implemented in pharmacology research. In our work the “dose-response curves” are equivalent to SPL-auditory response curves. This allowed us to maximise the information contained in each dataset and to quantitatively compare model parameters such as: Hill coefficient (steepness of slope), maximal asymptote (maximum σ ratio), and inflexion point (σ ratio at the steepest part of the slope). We did this using the *drm* function of *drc* package (Version 3.1-1, Ritz and Strebig, 2015).

We fitted four-part Log-Linear models to each individual Locust’s auditory response, with auditory nerve responses (σ ratio) as the dependent variable with treatment (control or starved/cold-reared) and SPL as the independent variables. Where relevant, a fixed effect was also calculated for Locust Sex, and the models for each individual Locust were adjusted accordingly. The t and p values are reported for each model parameter: Hill coefficient, maximal asymptote, and inflexion point - shown on each graph in Figures 5 and 6. The equation of the four-parameter log-linear fits is:

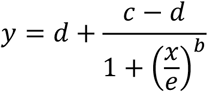

Where *Y* is the σ ratio, *b* is the slope at the inflexion point, *c* is the lower asymptote, *d* is the higher asymptote, *e* is the SPL (or *X* value) producing a response halfway between *b* and *c*.

To test whether the factor of treatment (starved or cold-reared condition or control and metabolic rate) significantly affected auditory nerve response we compared the above model to a model in which treatment was omitted as an independent variable, using the *anova* function (Ritz et al., 2015). This gave an F statistic labelled on each graph in Figures 5 and 6. Cohen’s d was then used to calculate effect size difference at the most different dB SPL. This dB SPL was then used in Fig.4C.

## Acknowledgements

We thanks Celia Hansen and Ramesh Patel for growing wheat, Chris Fisher for technical support, James Pinchin for help with analysis of hook electrode data for Figure 2, Tom Matheson for help with wheat watering,

## Author contributions

TA collected and analysed hook electrode data for figures 4, 5 and 6 and composed figures 1,4,5,6. TA extracted RNA for figure 7. AB collected and analysed hook electrode data for figure 2. CT, CL and TA analysed transcriptomic data for figure 7 and CT composed figure 7. BW collected and analysed metabolic assays for figures 1, 4, 5, 6 collected and analysed patch-clamp data for figure 3. BW, CT and TA wrote the manuscript.

## Competing interest

The authors declare no competing interests

## Material and Correspondence

## Notes

### Competing Interest Statement

The authors have declared no competing interest.

